# dismo: building and analysing discrete spatial ODE models

**DOI:** 10.1101/2023.10.17.562679

**Authors:** Marvin van Aalst, Oliver Ebenhöh

## Abstract

**Summary:** `dismo` is a Python package for building and analysing discrete spatial models based on ordinary differential equations. Its primary purpose is to allow arbitrarily complex internal and transport processes to easily be mapped over multiple different regular grids. For this it features one, two and threedimensional layouts, with standard and non-standard (e.g. hexagonal or triangular) grids.

**Availability and implementation:** https://gitlab.com/qtb-hhu/dismo

## Introduction

Complex multicellular organisms depend on communication between cells. This includes exchange of information, for example by hormone signalling, and the transport of metabolites between cells and organs. Many mathematical models of biological processes, including metabolism, signalling, and ecosystem dynamics, ignore this spatial dimension and describe the system dynamics by coupled ordinary differential equations. The classical approach to model spatial dynamics is by using partial differential equations (PDEs). Famous examples include reaction-diffusion systems, which are able to describe diverse phenomena, such as the formation of patterns or the dispersion of animals in an ecosystem [1, 2]. However, the numerical treatment of PDEs poses various challenges. Simulation requires a careful definition of the boundary conditions, and the spatial discretisation must fulfil certain criteria in order to guarantee convergence. Moreover, they are computationally very expensive, especially when the number of dynamic variables becomes large. However, when describing multicellular systems, the dynamics within each individual cell can be often considered to not be limited by diffusion, and thus can be approximated to be well-mixed. If this assumption is fulfilled, the dynamics can be described as the interaction of discrete entities, which may correspond to the single cells. As a consequence, in such a case the dynamics can be described using a system of ordinary differential equations (ODEs), in which the intracellular processes and the exchange processes between cells form the basis of the coupled equations. In this way, the spatial discretisation naturally reflects the biological system under investigation, and, especially when different cell types are involved, appears more straightforward than in a pure PDE-based approach.

While there are excellent tools for solving PDEs in Python, such as py-pde, to the best of our knowledge there exist no tools to create and analyse discrete spatial ODE models [3]. Here, we present dismo, a Python package for implementing and solving discrete spatial systems of ODEs. Our package provides methods to automatically create the complete set of coupled ODEs, based on the user-defined grid and the ODEs describing the processes within, and transport between the individual cells. Our package offers numerous regular pre-defined one-, twoand threedimensional grids, and supports the definition of different cell types, which can each have their specific intracellular dynamics. In this application note, we describe the functionality of dismo and demonstrate its capabilities for a spatially resolved model of sugar transport in a photosynthetic leaf.

**Figure 1.**
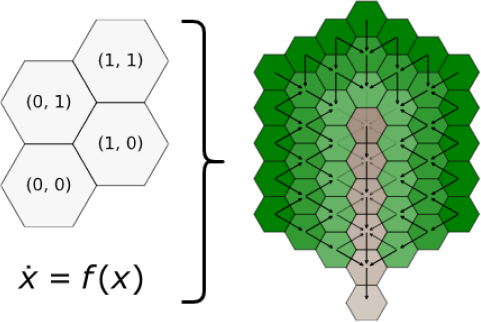
Schematic representation of a dismo model. The left-hand-side depicts that every model consists of a grid, which can be one-, two- or threedimensional, and one or multiple ordinary differential equations for one or multiple state variables, that either describe the dynamics within a given cell, or transport processes between cells in that grid. The right-hand-side depicts the flux of sugars in a plant leaf model.

## Implementation and results

We build `dismo` using best object-oriented programming (OOP) practices, utilizing composition, separation of concerns and abstract base classes [4]. For example, coordinates and their respective behaviour are separated from the general grid implementation, such that it is easy to implement new grids. Further, we utilized composition instead of inheritance to supply different grids for the model instances. This way it is easy to interchange parts and expand both the model and grid types. Construction and analysis of models, as well as how to subtype a model class are all described in detail in accompanying jupyter notebooks [5].

Common analysis methods like time course simulation and visualization are implemented as ready-touse functions and utilize well-known packages from the scientific Python landscape, like `NumPy, pandas, Matplotlib` as well as the `assimulo` wrapper around the `sundials` solver suite [6, 7, 8, 9, 10].

As modelling sugar transport inside of plant leaves was the motivation behind building `dismo`, the package also includes different types of plant leaf models, which build up in complexity. First we supply mesophyll models, which consist of a single variable for sucrose as well as a single mesophyll cell type and incorporate both a saturating photosynthesis function and a passive diffusion process between mesophyll cells. The mesophyll model is then expanded by supplying an additional vein cell type, which transports sucrose more rapidly and provides an outflux out the leaf. For this, passive transport processes between vein cells as well as an active (as in one-sided) transport of sucrose from the mesophyll cells into the vein cells are added to the model description. Finally, the stomata models further extend the model by both a new stomatal celltype and a second variable for CO2. The idea here is that CO2 only enters stomatal cells, which can then transport this CO2 into mesophyll cells. Vein cells are assumed not to contain any CO2. In this model the mesophyll photosynthesis function is now dependent on the CO2 concentration in that particular cell and thus more dynamic and subject to the placement of stomatal cells in the leaf.

## Conclusion

`dismo` is a new Python package developed for building and analysing multi-variate, discrete spatial models based on ODEs. It thus differs from packages focused on continuous PDEs, enabling spatial discretisation that naturally reflects the biological system under investigation, making model construction and analysis more straight-forward than in a PDE-based approach.

